# Continuous structural neuroplasticity during motor learning - a diffusion MRI study

**DOI:** 10.1101/2024.01.09.574830

**Authors:** Naama Friedman, Cfir Malovani, Inbar Perets, Etai Kenin, Michal Bernstein-Eliav, Ido Tavor

## Abstract

How does our brain transform when we encounter a new task? To fully answer this question, comparing brain states before and after learning may not be enough, but rather an on-going, continuous monitoring of brain changes during learning is required. While such continuous examinations of functional learning-induced changes are widely available using functional magnetic resonance imaging (fMRI), a continuous investigation of microstructural brain modifications during learning is yet to be reported.

Here, we continuously acquire diffusion MRI images during task performance. We then compute the mean diffusivity (MD) using a sliding-window approach, resulting in a continuous measure of microstructural changes throughout learning. We demonstrate the utility of this method on a motor sequence learning (finger tapping) task (n=58). MD decrease was detected in task-related brain regions, including the parahippocampal gyrus, hippocampus, inferior temporal gyrus, and cerebellum. Analysis of the temporal patterns of decrease revealed a rapid MD reduction in the right temporal gyrus after 11 minutes of learning, with additional decrease in the right parahippocampal gyrus and left cerebellum after 22 minutes. We further computed “neuroplasticity networks” of brain areas showing similar change patterns and detected similarities between these networks and canonical functional connectivity networks.

Our findings offer novel insights on the spatio-temporal dynamics of microstructural neuroplasticity by demonstrating continuous modifications during the encoding phase of learning itself, rather than comparing pre- and post-learning states.

## 1. Introduction

Learning takes time. While the development of new skills can occur over extended periods of practice and experience (Green and Bavelier, 2008; Knight et al., 2017), learning-related neural modifications are evident already at the very first minutes of encoding (Brodt et al., 2018; Sagi et al., 2012; Tavor et al., 2020). To fully understand the neural underpinnings of learning it is crucial to consider its continuously evolving nature.

Numerous studies have explored the structural and functional alterations that accompany learning using structural and functional magnetic resonance imaging (MRI), respectively. Changes in cortical thickness and gray matter volume have been reported (e.g., Koch et al., 2016; Legault et al., 2019), as well as changes in white matter structures (e.g., Hofstetter et al., 2013; Sampaio-Baptista et al., 2013; for a review see Sampaio-Baptista and Johansen-Berg, 2017). Functional brain changes have also been detected (e.g., following motor skill acquisition, Ungerleider et al., 2002; or musical training, (Luo et al., 2012), mainly in learning-related cortical areas, but more recently also in the cerebral white matter (Frizzell et al., 2022, 2020; Ji et al., 2023).

While structural changes have been so far described primarily on a macro-scale level, over the last decade diffusion-weighted MRI (dMRI) has been employed to study *microstructural* neuroplasticity in gray matter structures as well (Brodt et al., 2018; Hofstetter et al., 2017; Jacobacci et al., 2020; Keller and Just, 2016; Sagi et al., 2012; Tavor et al., 2020, 2013; Villemonteix et al., 2023). Crucially, this recently emerging line of investigation differs from previous research in that it focuses on structural rather than functional changes and on a *micro*-rather than macro-scale level. Additionally, while dMRI has been traditionally used to study white matter structures, here it is employed to investigate gray matter plasticity.

Diffusion MRI measures the translational displacement of water molecules (Bihan and Warach, 1995). This allows to construct a mathematical representation, the diffusion tensor (Basser et al., 1994), from which it is possible to calculate quantitative information about the tissue microstructure as summation indices, such as the mean diffusivity (MD). Learning-induced decrease in MD has been reported (Hofstetter et al., 2017; Sagi et al., 2012; Sampaio-Baptista et al., 2013; Taubert et al., 2010; Tavor et al., 2020, 2013), possibly reflecting changes in the extracellular matrix (Van Der Toorn et al., 1996), formation of synapses and dendrites (Toni et al., 1999), dendritic remodelling, axonal sprouting and pruning (Kays et al., 2012), neurogenesis (Knoth et al., 2010; Maćkowiak et al., 2002; Seib and Martin-Villalba, 2015), astrocyte remodelling (Assaf, 2018; Blumenfeld-Katzir et al., 2011; Sagi et al., 2012) or other modifications in several types of glial cells (Fields et al., 2014; Weston et al., 2015). MD may be therefore used as an anatomical biomarker for neuroplasticity.

Neuroplasticity studies using dMRI have been focusing on various timeframes of brain changes, ranging from a few months to a few days (Thomas and Baker, 2013) and even just a few hours of training across different cognitive domains, including spatial navigation (Keller and Just, 2016; Sagi et al., 2012; Tavor et al., 2013; Villemonteix et al., 2023), motor sequence training (Jacobacci et al., 2020; Tavor et al., 2020), language (Hofstetter et al., 2017) and associative memory (Brodt et al., 2018). Notably, these studies have investigated the differences in brain microstructure between two discrete timepoints: before and after a learning routine. However, the temporal dynamics of the learning process itself, and what transpires within the brain during that time, have not been fully explored yet.

This study aims to monitor the continuous microstructural changes occurring in the brain during the learning process itself. To achieve this, we computed a continuous MD measurement and used it to identify temporal patterns of neuroplasticity across the brain, to detect the exact changepoint in MD during learning and to define ‘neuroplasticity networks’ of brain regions that share common microstructural modification patterns.

## 2. Methods

### 2.1 Participants

Sixty-two right-handed healthy volunteers with no history of neurological disease, psychiatric disorders, drug or alcohol abuse, or use of neuropsychiatric medication were recruited. Four participants were excluded due to technical issues during the scan or excessive movement, resulting in a final sample of fifty-eight participants (30 females, mean age = 27.66, SD = 4.4). Participants were randomly assigned to either a learning group (29 participant, 14 females, mean age = 28.47, SD = 4.26) or a control group (29 participant, 16 females, mean age = 26.5, SD = 4.37; with no significant age or gender differences between groups). The experimental protocol was approved by the Institutional Review Board of the Sheba Medical Center and all participants signed an informed consent form.

### 2.2 Experimental design

Participants underwent an MRI session which included a dMRI scan and several structural scans. During the dMRI scan, participants in the learning group performed a motor sequence learning task (see details below) while participants in the control group were instructed to focus on a fixation cross and not perform any explicit task. MRI acquisition was carried out in The Alfredo Federico Strauss Center for Computational Neuroimaging at Tel Aviv University.

### 2.3 MRI acquisition

Scans were acquired on a 3T whole-body MRI system (Siemens Magnetom Prisma) equipped with a 64-channel head coil.

Diffusion-weighted images (DWI) were acquired with a spin-echo diffusion-weighted, echo-planar imaging sequences with up to 68 axial slices (whole-brain coverage) and a resolution of 2×2×2 mm^3^, with a TR/TE = 3,500/59.4ms, multiband acceleration factor = 2. Diffusion parameters were: Δ/δ= 28/10ms, b-value of 1,000s/mm^2^ were taken at 589 gradient directions with additional 33 non-diffusion weighted (b_0_) images (spread evenly across the scan duration, such that in each MD calculation, the nearest B_0_ image was used as a reference to assess diffusion-related decay of the MR signal for each tensor calculation). The total dMRI scan duration was 36:28 minutes. In addition, five b_0_ images and one b_1000_ image were acquired with a reversed phase-encoding direction to correct for susceptibility-induced distortions as described in section 2.6.

T1-weighted images were acquired with a magnetization prepared rapid gradient echo (MPRAGE) sequence with up to 176 axial slices (whole-brain coverage), TR/TE=2,400/2.98ms, resolution of 0.9×0.9×0.9 mm^3^, scan time of 4:30 minutes. T2-weighted images were acquired with up to 176 slices (whole-brain coverage), TR/TE=3,200/554, resolution of 0.9×0.9×0.9 mm^3^, scan time of 5 minutes. In addition, fluid attenuated inversion recovery (FLAIR) images (TR/TE/TI = 8,000/81/2,370) were acquired for radiological screening.

### 2.4 Motor sequence learning task

Participants in the learning group performed a finger tapping task (first introduced by Karni et al., 1995). The task consisted of 24 trials of 60 seconds, each followed by a 25-second rest period, for a total duration of 33:35 minutes, with additional 2:53 minutes of rest at the end of the task. During task trials, participants were instructed to repeat a 5-digit sequence (2-4-1-3-2) using their left (non-dominant) hand as correctly and quickly as possible, while the sequence is constantly presented on the screen to eliminate executive memory differences. Rest periods were added to allow for offline gain learning (Bönstrup et al., 2019; Jacobacci et al., 2020). Finally, participants performed an additional task trial while in the scanner, but with no scan being performed, serving as a behavioral test.

### 2.5 Behavioral assessment

Two (non-independent) measurements of task performance were calculated: (1) accuracy: the total amount of correct sequences completed, calculated as the number of correct sequences a participant performed within a specific trial, and (2) speed: the mean duration of a successful key press sequence, calculated as the average duration of correct sequences in a specific trial (sequence duration being the time passed between pressing the first and last keys of a sequence). Both measurements were computed for each trial and for the test trial. Paired t-tests were performed for each of these measurements, between the first and last trials, and the test trial.

### 2.6 dMRI data analysis

dMRI images were corrected for head motion, susceptibility-induced distortions, and eddy currents-induced distortions, as well as registered to the T1-weighted image using the minimal preprocessing pipeline proposed by the Human Connectome

Project (HCP) (Glasser et al., 2013; Sotiropoulos et al., 2013) including the FMRIB Software Library (FSL) TOPUP and EDDY tools (Andersson et al., 2003; Andersson and Sotiropoulos, 2016). For optimized registration to the Montreal Neurological Institute (MNI) standard space we applied a tissue-probability based registration routine, as recently described by Malovani et al., 2021. Data were spatially smoothed with a gaussian kernel with a full width at half maximum (FWHM) of 5 mm using FSL.

To obtain a continuous measurement of microstructural changes during learning, we calculated the diffusion tensor by applying a sliding window approach: We used a window length of 13 TRs (12 DWI images plus one non-diffusion weighted (b_0_) image), and a window offset of a single TR as described in Figure 1a. To determine the optimal window length, we used data from three different sources, using three different diffusion protocols: (1) Four participants who were excluded from the present study due to technical issues or excessive movement. These participants were scanned with the exact same diffusion protocol as the remaining participants in this study (see section 2.3 above). (2) Ten participants who were scanned in the same MRI scanner but using a different dMRI protocol, containing 64 different gradient directions at b=1000 s/mm^2^. (3) Five HCP participants (Van Essen et al., 2012) who underwent a dMRI scan containing 90 different gradient directions at b=1000 s/mm^2^. We calculated the MD from an increasing number of directions, ranging from 1 to 61 (to allow for an adequate number of possible permutations for the shortest dataset (number 2), or 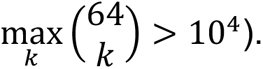 To calculate stability, we created 1000 permutations for each number of directions, and calculated MD each time using a different set of directions. Stability was defined by the standard deviation (SD) of the average MD in the total volume of the gray matter (see Supplementary Figure S1-3 panel A) and in four specific areas: the postcentral gyrus, the fusiform gyrus, the caudate nucleus, and a part of the cerebellum, for each number of gradient directions (see Supplementary Figure S1-3 panel C). The stability of the MD measurement increased with the number of directions used, with more than 11 directions resulting in near-zero values of SD across permutations (see Supplementary Figure S1-3 panel B). Therefore, we concluded that 12 gradient directions are sufficient for a stable MD calculation. For each window of 12 DWI images, we added the nearest b_0_ image for MD calculation. Window offset was set to 1 TR to maximize the number of MD timepoints. Considering these window length and offsets, and a TR of 3.5 seconds, an MD timepoint was calculated based on 42 seconds of data and the difference between MD timepoints was 3.5 seconds. MD values were calculated along the timeseries of the DWI scan, resulting in a continuous measurement of MD during learning (Figure 1). To this continuous curve we applied a lowpass filter with a cut-off frequency of 0.0015Hz to remove high frequencies unrelated to the MD decrease trend (in Supplementary Figure S5 we present the unfiltered continuous curve). All analyses were performed in MATLAB (2020b, Natick, Massachusetts: The MathWorks Inc). Our code used for continuous MD calculation is available at: https://github.com/naamaf/Continuous_DTI.

**Figure 1.**
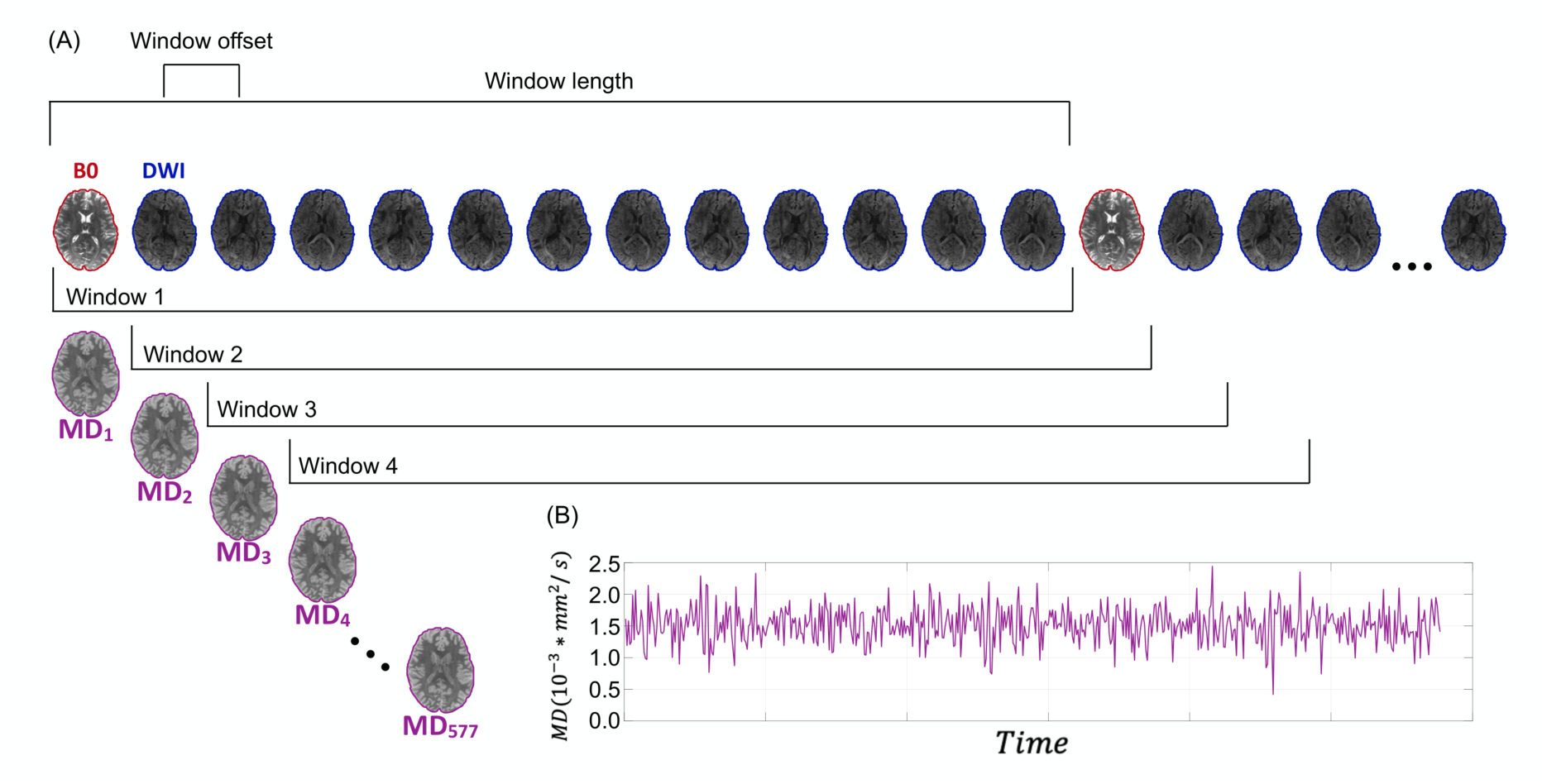
Continuous MD using a sliding window approach. **(A)** An illustration of a sliding window over multiple diffusion weighted images (DWI) volumes. Window length was set to 13 TRs (12 DWI images in different gradient directions + the nearest b_0_ volume) and the window offset was one TR. **(B)** For each window, an MD value in each voxel was calculated to create a continuous measurement of microstructural properties.

In addition to this continuous characterization of MD changes, we also calculated MD “pre” and “post” learning, as the mean MD in the first and last ten MD timepoints in the continuous MD curve, respectively (see Supplementary Figure S4 for the same analysis with 1, 5 or 15 MD timepoints).

To further explore the evolving pattern of MD changes during learning before inspecting the full continuous change in time, we investigated discrete timepoints along the continuous MD curve shown in Figure 1B. Specifically, we computed the percent change in MD between the “pre-learning” timepoint (as described above) and three additional timepoints: around 11 minutes into the learning process, around 22 minutes into the learning process, and at the end of the scan, after learning is completed, which is identical to the “pre”/”post” comparison described above. The timepoints at 11 and 22 minutes were chosen as two middle points during the learning process. Each of these additional points was calculated as the mean of 10 MD timepoints, starting at 11 or 22 minutes into learning. Since an MD timepoint lasted 42 seconds and was separated by 3.5 seconds from the next one, 10 MD timepoints lasted 42 + (3.5 ∗ 9) = 73.5 seconds (i.e., “11 minutes into learning” was the mean MD from the 11^th^ minute to the 12 minute and 13.5 seconds, and “22 minutes into learning” was the mean MD from the 22^nd^ minute to 23 and 13.5 seconds).

### 2.7 Statistical analysis

To detect clusters of MD decrease, that were later used for investigation of continuous modifications during the learning process itself, we first performed a voxel-wise two-way ANOVA of MD timepoint (before / after learning) X group (learning/control) using FSL’s PALM tool with 1000 permutations (Winkler et al., 2014). Only gray matter voxels were included, and a threshold-free cluster enhancement (Smith and Nichols, 2009) was utilized. *P*-values were FDR corrected for multiple comparisons. Clusters with over 50 voxels that showed an interaction effect (i.e., MD decrease in the learning but not the control group) were included in following stages of analysis.

For each of these clusters, we computed the “pre” and “post” learning MD as described above, as well as the differences between the “pre-learning” MD and the additional timepoints throughout the learning process, averaged within the learning or control groups. Paired t-tests were performed between the “pre-learning” MD and the three subsequent timepoints (11 minutes, 22 minutes, and post-learning).

Next, to obtain a continuous characterization of MD change, we averaged MD values per MD timepoint within each cluster across voxels and across participants for each group independently, resulting in two (for the learning and control groups) distinct MD timeseries per cluster.

### 2.8 Changepoint detection analysis

To further compare between the patterns of MD change across clusters, we performed a changepoint analysis (Killick et al., 2012) per timeseries to detect a single specific timepoint in which the greatest change in slope has occurred. The changepoint for each cluster was calculated per participant and averaged per group. To achieve a more detailed understanding of the spatial pattern of temporal changes, changepoint analysis was additionally used to examine the change in slope separately for each voxel within a cluster.

### 2.8 Networks analysis

We next examined the similarities between microstructural plasticity patterns across brain areas. Gray matter was parcelled into 273 areas (Brainnetomme atlas; Fan et al., 2016). We calculated the individual continuous MD change averaged across voxels per parcel and computed Pearson’s correlations between all region pairs. Individual correlation matrices were then averaged across participants. We applied a hierarchical clustering on the averaged correlation matrices using Python library SciPy (Virtanen et al., 2020) to detect “neuroplasticity networks”, i.e., networks of brain areas that share similar temporal patterns of MD change during learning. This analysis resulted in a division of the brain into five “networks” (the optimal number of clusters was determined by the silhouette value (Rousseeuw, 1987). To examine the correspondence between these “neuroplasticity networks” and known functional networks, we computed the Dice index between each of the five “neuroplasticity” and the seven canonical resting-state functional connectivity networks (Yeo et al., 2011). Since the Yeo networks do not include sub-cortical regions or the cerebellum, those areas were excluded for the purpose of this comparison. For the same analysis with an alternative parcellation, using the Glasser atlas (Glasser et al., 2016), see Supplementary Figure S7-9.

## 3. Results

### 3.1 Behavioural results

Participants showed successful learning in both accuracy and speed (Figure 2). Comparing the first and last trials of learning, the mean improvement in speed was 42.9% (SE = 2.8%) and the mean improvement in accuracy was 67.2% (SE = 11.2%).

**Figure 2.**
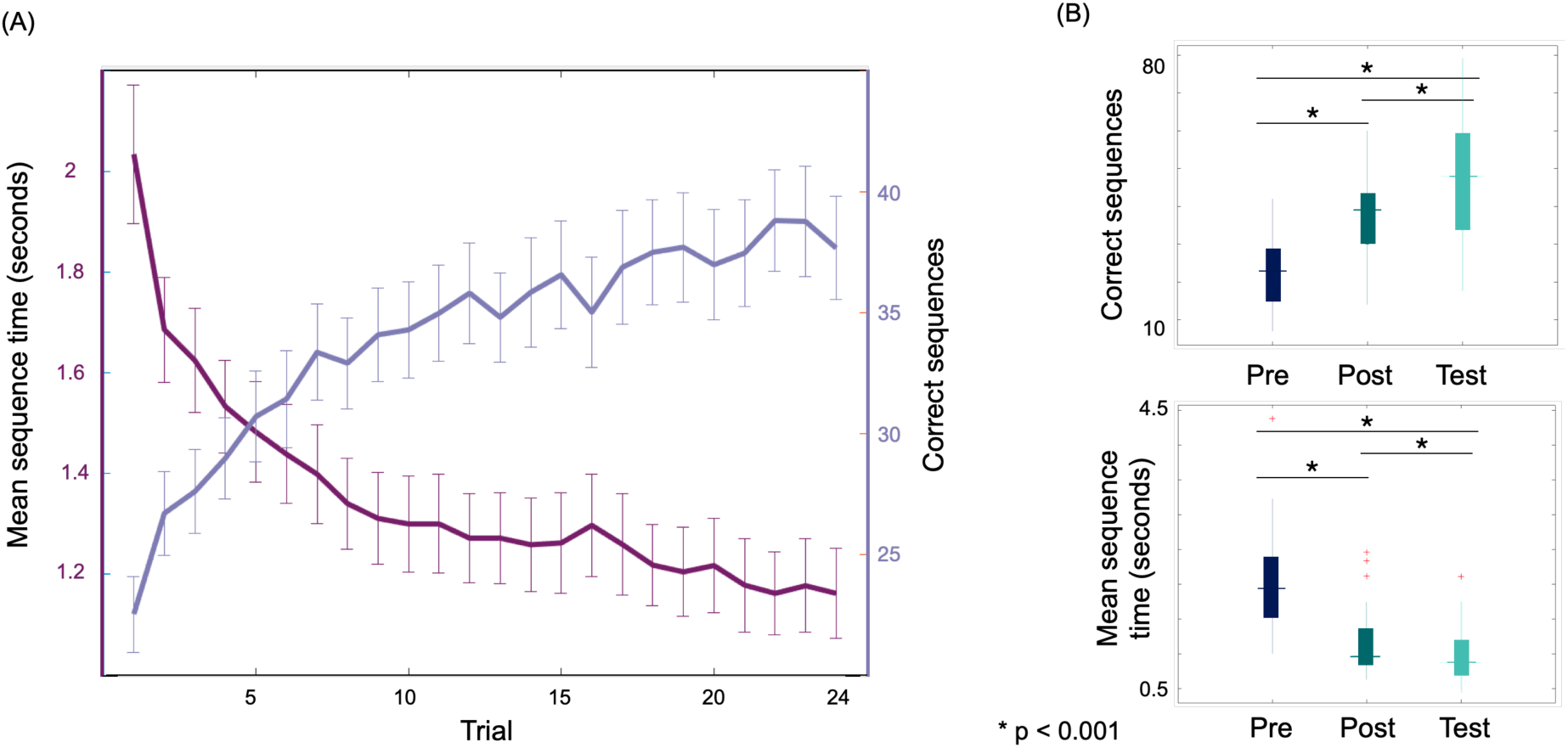
Behavioral effects of motor sequence learning. **(A)** Averaged behavioral scores for 29 participants who performed a finger tapping task for 33:35 minutes (24 trials). The figure shows, on the right y axis, the mean sequence duration per trial, and on the left y axis the mean number of correct sequences per trial. Error bars represent the standard error per trial. **(B)** Differences in performance between the first, last and test trials (paired t-test, *p* < 0.001, FDR corrected).

In the test trial, performed after a 12-minute break (inside the scanner), the improvement in speed was 52.3% (SE = 2.2%) and in accuracy, 108.55% (SE =14.7%) compared to the first trial. Differences between the first, last and test trials were significant for both measurements (paired t-test, *p* < 0.001, FDR corrected).

### 3.2 Microstructural neuroplasticity at discrete timepoints throughout learning

In a voxel-wise two-way ANOVA of MD timepoint (before/after learning) X group (learning/control) we found a 1-4% MD decrease in the right parahippocampal gyrus (PHG), the right hippocampus, the right inferior temporal gyrus (ITG) and the cerebellum (Figure 3A and B). To further explore the evolving pattern of MD decrease during learning, we examined additional timepoints throughout the learning process. Each panel in Figure 3C includes, therefore, three bars, depicting MD change between the “pre-learning” baseline and the following timepoints: around 11 minutes into the learning process; around 22 minutes into the learning process; and after learning is completed, which is identical to the bars shown in Figure 3B. A rapid reduction in MD within the right temporal gyrus was observed after 11 minutes of learning, extending consistently until task’s completion. A decrease in MD was also observed after 22 minutes of learning in the right PHG and the left cerebellum.

**Figure 3.**
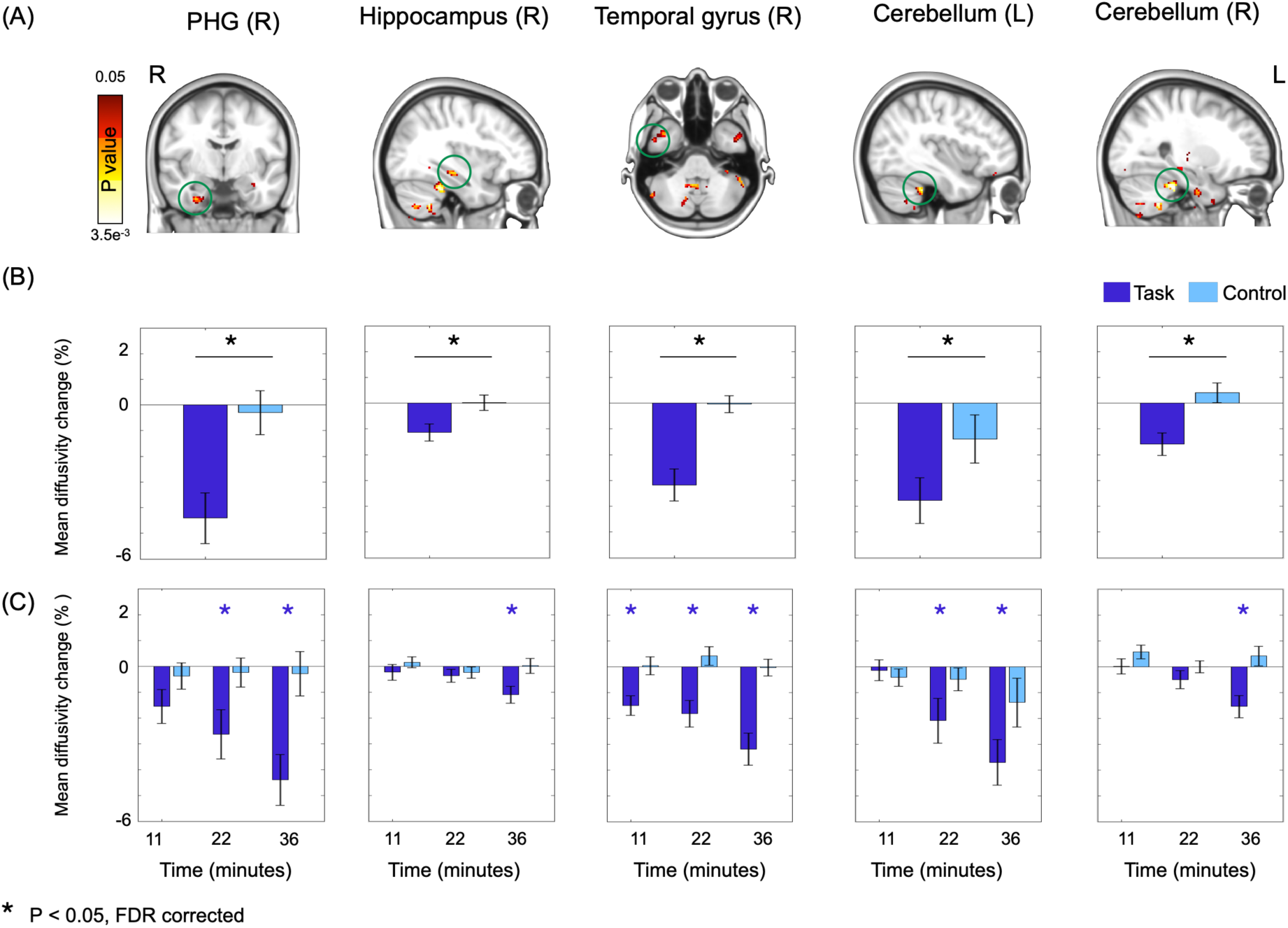
Microstructural neuroplasticity at discrete timepoints throughout motor sequence learning. **(A)** We first examined the pre/post learning effect following training: A 2 (learning vs. control groups) by 2 (pre/post) ANOVA revealed a significant interaction effect. Clusters of MD decrease following learning were found in the right parahippocampal gyrus (PHG), the right hippocampus, the right inferior temporal gyrus, and the cerebellum. **(B)** MD change in the clusters displayed in A, for the learning (dark blue) and control groups (light blue). **(C)** A three-point temporal course of MD change in the same clusters. Paired t-tests were performed between each timepoint and the pre-learning timepoint, asterisks denote a significant MD decrease in the learning group. Note that the rightmost bars in each plot in panel 3C are identical to those presented in panel 3B. Error bars represent the standard error.

### 3.3 Continuous training-induced microstructural neuroplasticity

The continuous MD change during learning is presented in Figure 4. Notably, within all clusters, the reduction in MD manifested gradually during learning; however, distinct variations were detected across regions.

**Figure 4.**
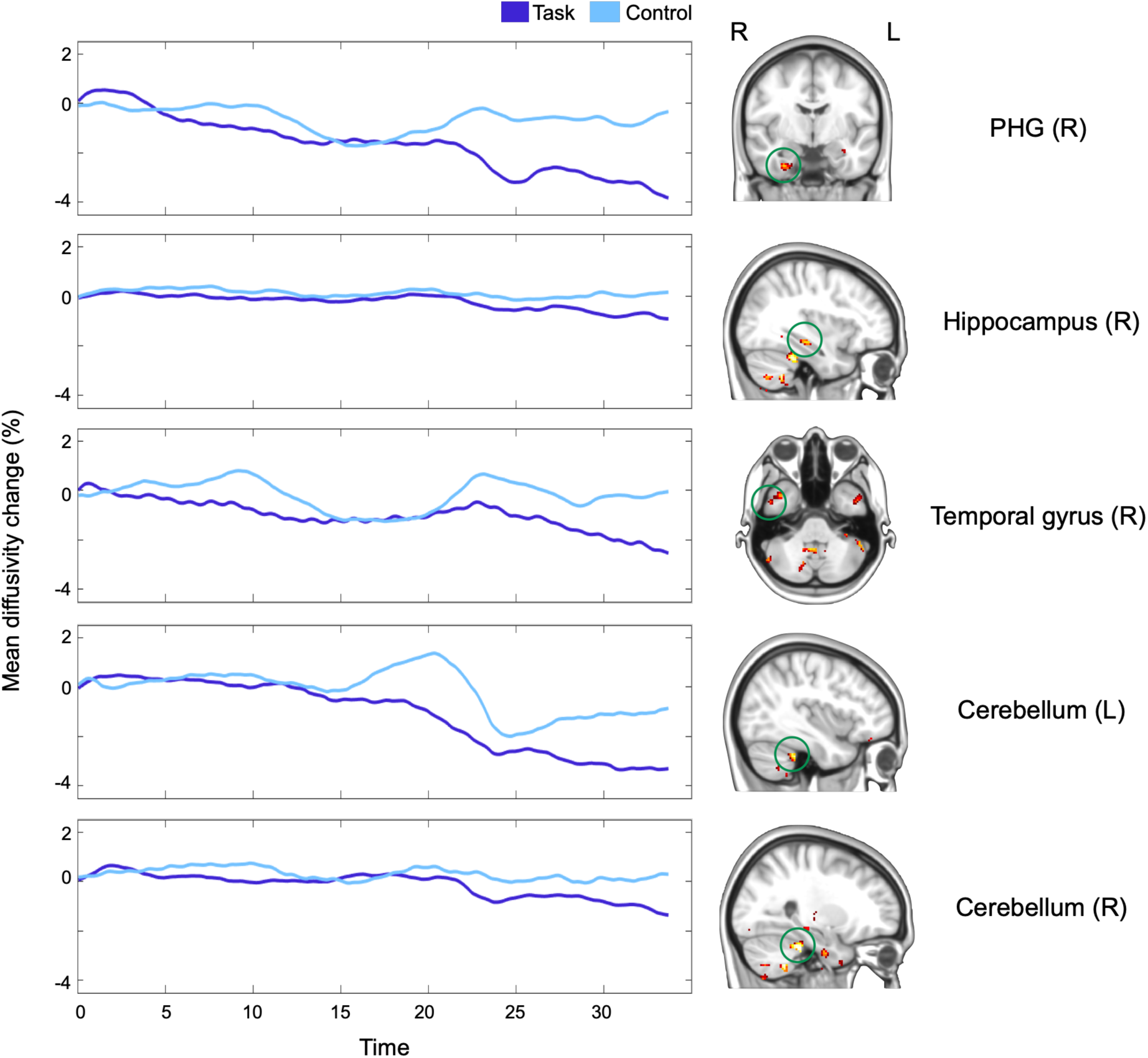
Continuous MD changes during learning. MD timeseries per cluster are shown after applying a lowpass filter of 0.0015 Hz (for unfiltered curves see Supplementary Figure S5). Curves were calculated individually and averaged across participants per group. While a decrease in MD was found in all areas for the learning group compared to the control group, the patterns of decrease differed across areas.

In Figure 5, we present the results of a changepoint analysis aimed to examine variations in the slope of MD decrease across clusters. Notably, in the parahippocampal gyrus we observed a relatively consistent slope throughout the learning process, while in the hippocampus and cerebellum we detected a significant changepoint occurring approximately 17 and 20 minutes into the learning task, respectively. During subsequent phases of learning, the MD decrease was considerably steeper compared to initial stages (Figure 5A).

**Figure 5.**
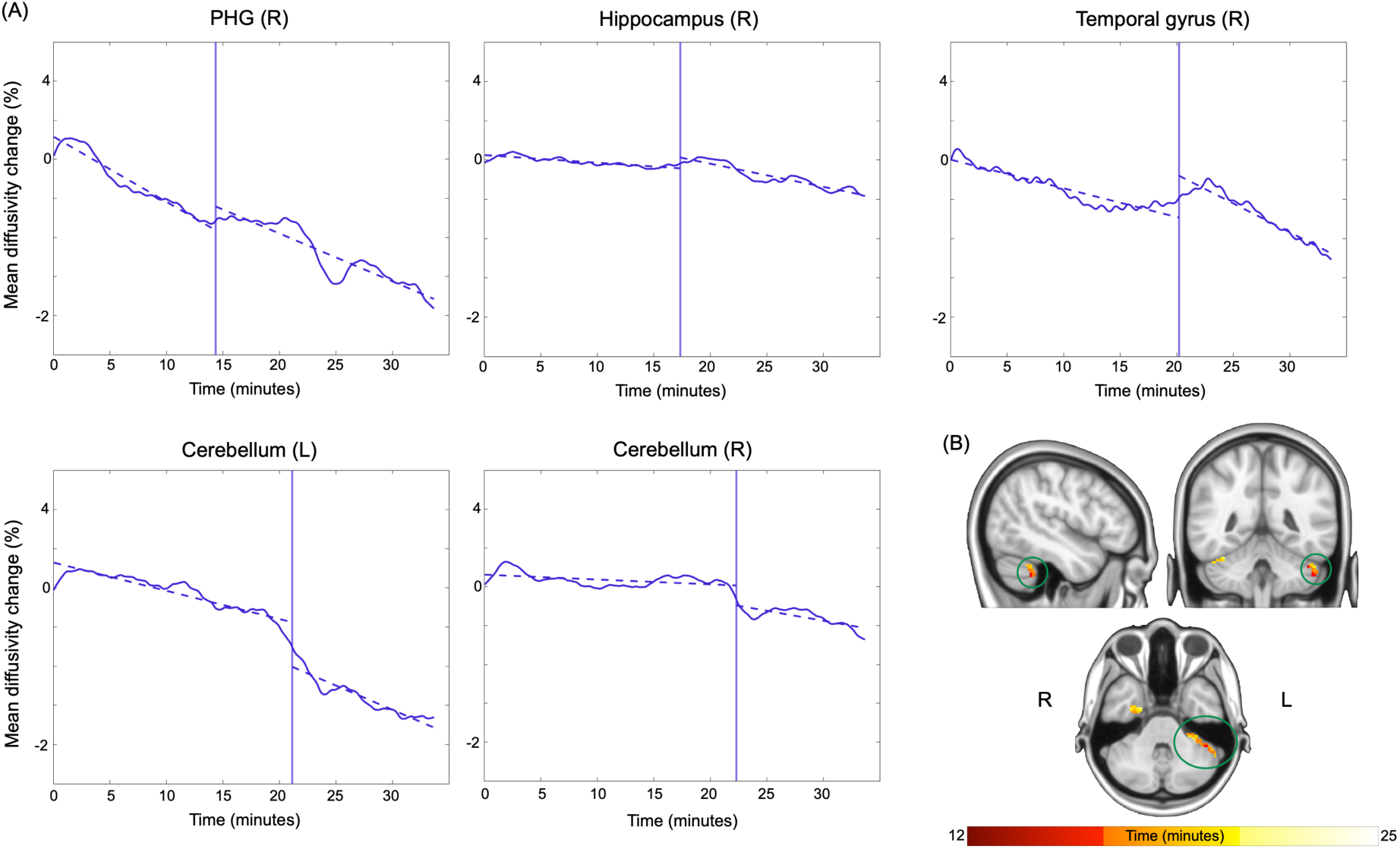
Changepoint detection analysis. **(A)** For each of the five clusters, a change in slope was determined as the maximum change between the slope of the two parts of the divided curve. The changepoint was calculated individually and averaged across participants and is presented on the average MD timeseries. **(B)** Voxelwise changepoint detection within the left cerebellum. Color-coding represents the time (minutes into the learning process) of changepoint for each voxel in the cerebellum.

It is worth highlighting that variations in the progression of MD decrease were not only evident between different brain regions but also within each area. For instance, in the cerebellum, we identified voxels displaying a change in slope around the 15-minute mark, while others exhibit this change at later timepoints (Figure 5B, see Supplementary Figure S6 for the voxelwise changepoint analysis results in other clusters).

### 3.4 Neuroplasticity networks

We next searched for “neuroplasticity networks”, reflecting a network-level organization of brain regions that undergo similar microstructural changes during learning. To examine the similarities in the patterns of MD decrease across brain regions, we first parcelled the brain into 273 areas based on the BNA atlas. We then calculated the continuous MD for each parcel across all learning participants and computed the Pearson’s correlations between each pair of parcels, resulting in a 273×273 correlation matrix. We then used hierarchical clustering to characterize five “neuroplasticity networks” (Figure 6).

**Figure 6.**
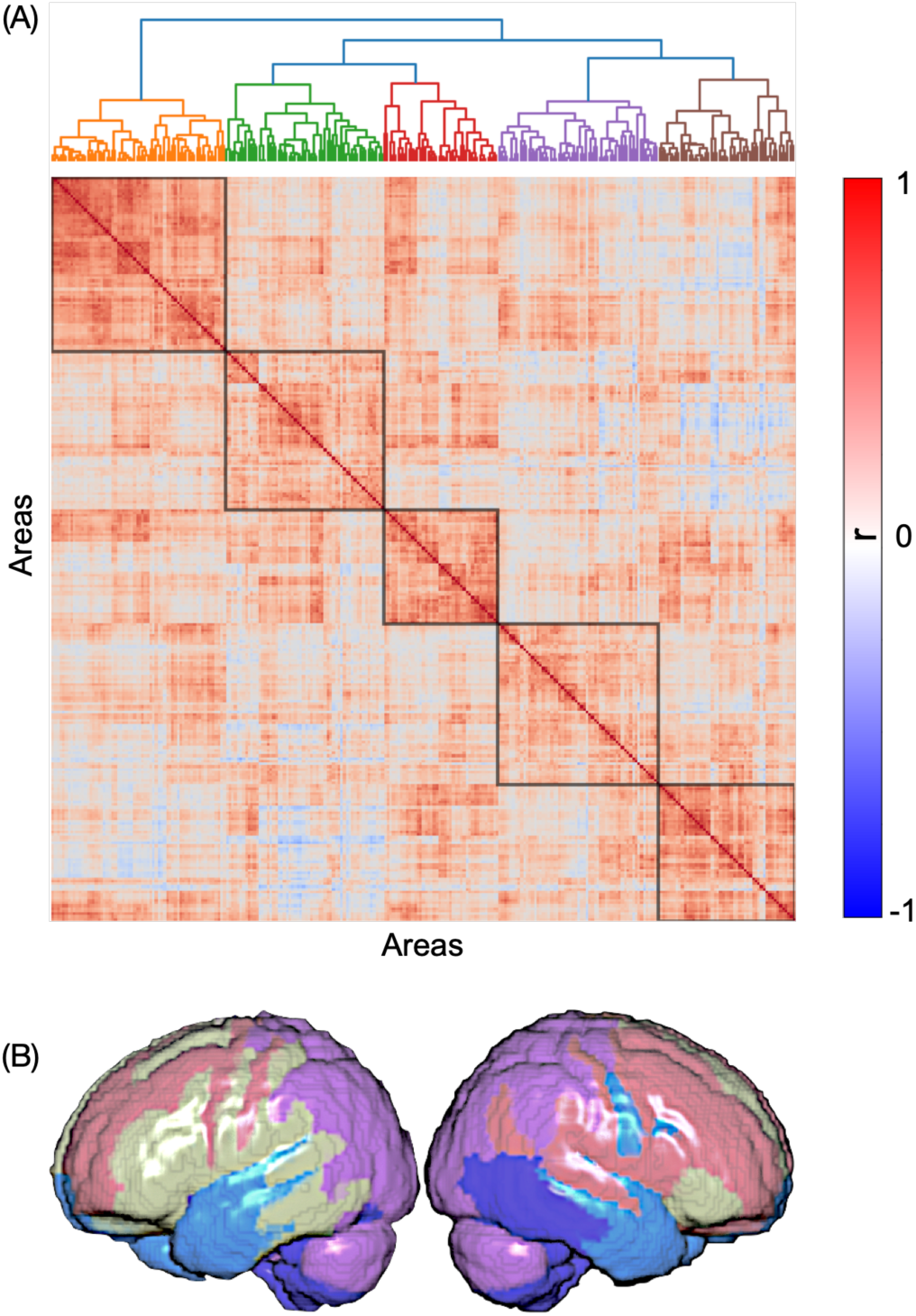
MD-change networks. **(A)** A matrix depicting the pairwise Pearson’s correlations between the patterns of MD decrease across 273 brain regions, ordered by a hierarchical clustering algorithm and parcelled to “MD-change networks” (black squares). **(B)** The five networks displayed on an inflated surface of the brain.

Finally, we examined the correspondence between these networks and canonical functional connectivity networks (Yeo et al., 2011) by computing the Dice index between each pair of networks (Figure 7). Similarities were detected between each of our “neuroplasticity networks” and either the limbic, the control or the visual networks.

**Figure 7.**
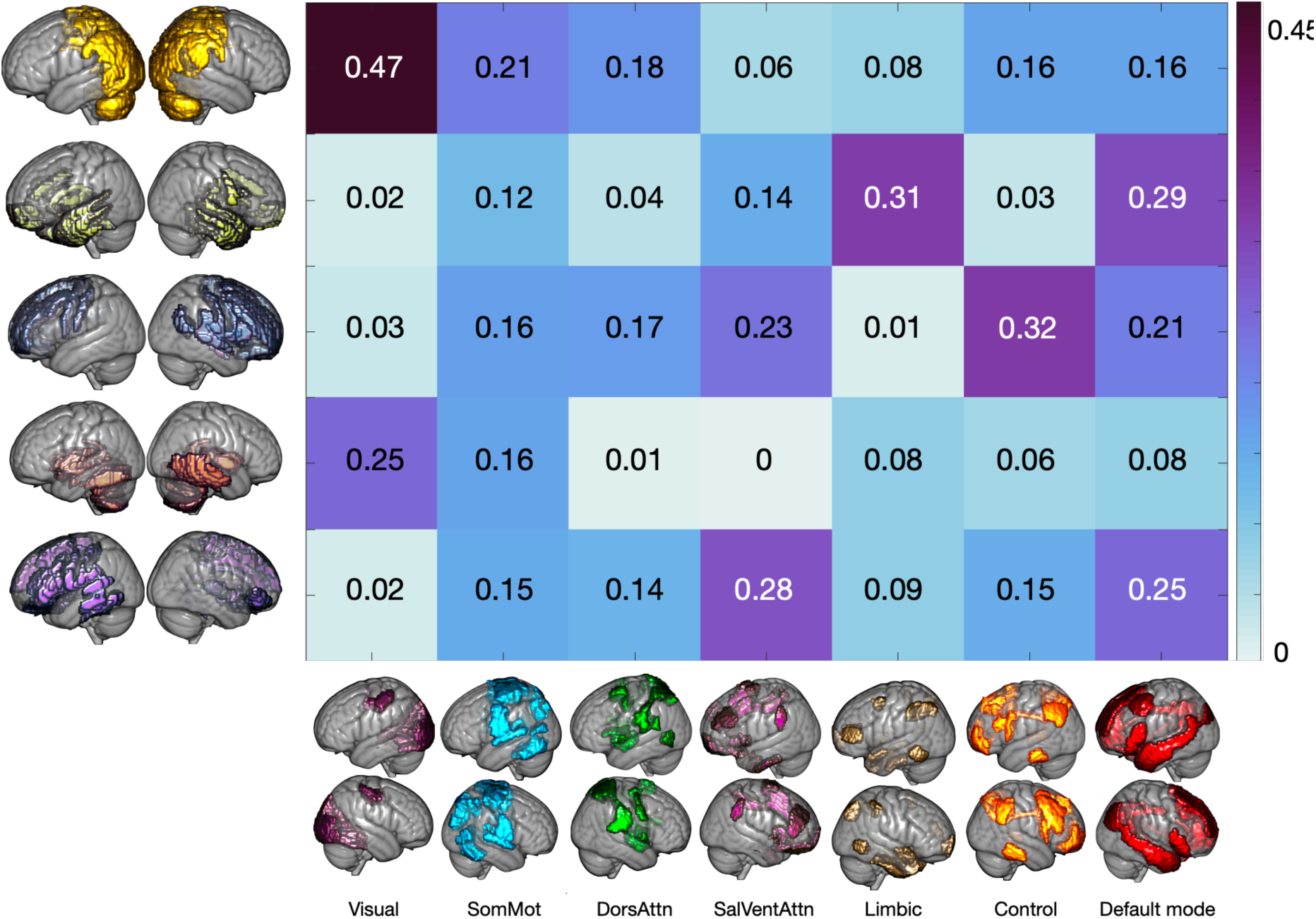
Similarities between MD-change networks and functional connectivity networks. Similarity matrix showing the Dice index between each of our five main neuroplasticity networks and Yeo’s seven connectivity networks. Highest Dice indices were found between the MD-change networks and the limbic network (Dice = 0.31), the control network (Dice = 0.32) and the visual network (Dice = 0.47).

## 4. Discussion

In this work we investigate continuous learning-induced microstructural remodelling of brain tissue, leveraging a unique diffusion MRI protocol to continuously assess mean diffusivity (MD) in gray matter throughout a motor learning routine. This protocol allowed us to detect not only learning-induced changes between discrete timepoints, as was done previously and also shown here in figure 3, but critically also the continuous change pattern throughout the entire learning process, as shown in figure 4. Different patterns of MD decrease were found in several learning-related brain areas, suggesting an involvement of these areas at the different stages of learning. Specifically, while the parahippocampal gyrus demonstrated a relatively consistent slope of MD decrease, the hippocampus and cerebellum showed a significant change in slope 17 to 20 minutes into the learning process. Furthermore, we were able to capture different change patterns across single voxels within a cluster, suggesting high sensitivity of our method to the spatio-temporal properties of learning-induced changes. Finally, based on similarities in the patterns of MD decrease across the brain we detected “neuroplasticity networks”, consisting of areas that undergo similar microstructural alterations during learning.

Our main goal was to examine where do signatures of neuroplasticity develop in the brain and how do they change over ongoing training. While the use of diffusion MRI to study short-term microstructural plasticity has gained increasing interest over the last decade (Brodt et al., 2018; Hofstetter et al., 2017; Jacobacci et al., 2020; Keller and Just, 2016; Sagi et al., 2012; Tavor et al., 2020, 2013; Villemonteix et al., 2023), these studies have all investigated the differences in brain structure between two discrete timepoints: before and after learning. This work is therefore the first to explore the temporal dynamics of microstructural changes as they occur during the encoding stage of the learning process itself.

Motor sequence learning requires a complex ensemble of distributed brain regions which are involved at different stages of learning (Dahms et al., 2020; Penhune and Steele, 2012; Tavor et al., 2020). Based on evidence from functional imaging studies (e.g., fMRI or PET) it has been suggested over 20 years ago that early stages of motor learning involve cortico-cerebellar and cortico-striatal mechanisms, whereas later stages of consolidation and retention engage the striatum, motor and parietal cortices (Toni et al., 1998; Doyon et al., 2003). Our results go beyond these classical models of learning by, first, describing microstructural rather than functional dynamics and second, by ‘zooming in’ into the very first minutes of motor learning, tracking continuous changes and detecting the exact moments in time when different brain areas join the process.

Of particular interest is the involvement of the hippocampus, which has been rarely reported in studies of functional plasticity following motor learning (Berlot et al., 2020; Coynel et al., 2010) but is in line with recent diffusion MRI studies showing an early engagement of the hippocampus during learning (Brodt et al., 2018; Jacobacci et al., 2020). Our findings may support a role of the hippocampus in short-term memory stabilization (Schapiro et al., 2019) or in connecting encoded items of different modalities over space or time (Buzsáki and Tingley, 2018).

While it is evident that the unique diffusion MRI protocol developed here is sensitive to dynamic, flexible brain modifications over short timescales, the biological underpinnings of such modifications are yet to be fully understood. A possible explanation is that learning-induced alterations in MD may reflect the rapid modification of astrocytes structures during and following learning (Assaf, 2018; Johansen-Berg et al., 2012; Sagi et al., 2012; Tavor et al., 2013).

The division to ‘neuroplasticity networks’ based on similarities of MD change patterns across brain regions offers further insights into the complex, multi-regional process that ultimately gives rise to learning and memory. These networks are not to be confused with the widely studied functional or structural networks, which reflect the temporal synchronization in brain activity across regions or the physical connections between them, respectively. Considering the accumulating evidence on a high correspondence between functional connectivity and brain activity (Gal et al., 2022; Smith et al., 2009; Tavor et al., 2016; Tik et al., 2023, 2021) and the recent suggestion that functional network architecture may relate brain activity and behaviour (Bernstein-Eliav and Tavor, 2022; Bijsterbosch et al., 2020), a network-organization approach may facilitate the study of neuroplasticity and its behavioural manifestation as well. Specifically, grouping areas by the patterns of microstructural changes may shed new light on the intricate mechanisms underlying skill acquisition.

An examination of the hierarchical clustering results reveals that the largest ‘neuroplasticity networks’ may roughly correspond with the functionally-defined limbic and control networks (Yeo et al., 2011), which is consistent with the known involvement of frontal, temporal and parietal cortices in motor sequence learning (Turesky et al., 2018; Witt et al., 2008). This correspondence is particularly intriguing as Yeo’s networks are defined based on the functional MRI signal during resting-state while our ‘neuroplasticity networks’ are driven from diffusion properties during task performance. It may therefore provide converging evidence for neuroplasticity mechanisms supported by brain areas within the control network, including the prefrontal, premotor, and parietal cortices which play a critical role in motor learning and execution. Interestingly, disrupted functional connections between areas in the control and limbic networks have also been associated with motor deficits in ageing (Michely et al., 2018), neurodegenerative disorders (Poston and Eidelberg, 2012), or following a stroke (Siegel et al., 2016).

This work introduces a novel method to continuously track diffusion properties throughout learning. Our results provide a promising proof-of-concept, yet further research is required to examine methodological aspects and optimize parameters such as window length and offset. It should also be considered that due to the nature of MD, which is calculated based on a series of DWI images acquired from various directions at intervals of a few seconds (depending on the repetition time, TR), the temporal resolution of our method is lower than that of functional imaging methods such as fMRI or even functional diffusion MRI (fdMRI) (Le Bihan, 2003; Nunes et al., 2021, 2019; Tsurugizawa et al., 2013). Therefore, further research is required to address the correspondence between these different imaging methods and integrate between them to achieve a more comprehensive picture on the temporal dynamics of learning-induced changes. Additionally, in this study we employed a relatively simple motor learning task; future work may apply our method to study higher-level cognitive tasks and complex learning routines. Insights on the timing and the network-level organization of learning-induced microstructural changes may extend previous findings on neuroplasticity following spatial navigation (Keller and Just, 2016; Sagi et al., 2012; Tavor et al., 2013; Villemonteix et al., 2023), language (Hofstetter et al., 2017) and associative memory tasks (Brodt et al., 2018).

In conclusion, we provide evidence for the utility of a continuous acquisition of diffusion MRI during task performance to explore the dynamic aspects of microstructural neuroplasticity. Rather than a single, post-learning ‘snapshot’ that indicates that changes have occurred at some, undetectable, point and pace, we show that different task-related brain areas demonstrate distinctive temporal patterns of learning-related modifications. The continuous characterization of neuroplasticity is therefore a promising approach, offering a temporally sensitive, network-level view on the orchestrated processes underlying learning and memory in the human brain.

## 5. Data and Code Availability

The code used for continuous MD calculation is available at: https://github.com/naamaf/Continuous_DTI. Raw MRI data cannot be shared due to data protection, but we provide tables of continuous MD values across parcels in the code repository.

## 6. Author Contributions

Naama Friedman: Conceptualization, Data collection and Curation, Methodology, Writing - Original Draft; Cfir Malovani: Formal analysis, Validation; Inbar Paretz: Data collection; Etai Kenin: Data collection; Michal Bernstein-Eliav: Writing – Review & Editing; Ido Tavor: Supervision, Conceptualization, Writing - Review & Editing.

