## Supplementary for "Continuous structural neuroplasticity during motor learning - a diffusion MRI study"

### Supplementary Information

Naama Friedman[1], Cfir Malovani[2], Inbar Perets[3], Etai Kenin[4], Michal Bernstein-Eliav[1], Ido Tavor[1,3,5]

[1]Faculty of Medical & Health Sciences, Tel Aviv University, Tel Aviv, Israel

[2]School of Electrical Engineering, Faculty of Engineering, Tel Aviv University, Tel Aviv, Israel

[3]Sagol School of Neuroscience, Tel Aviv University, Tel Aviv, Israel

[4]Department of Natural and Life Sciences, The Open University of Israel, Israel

[5]Strauss Center for Computational Neuroimaging, Tel Aviv University, Tel Aviv, Israel

**Corresponding Author:**

Dr. Ido Tavor; ORCID ID https://orcid.org/0000-0002-9117-4449

Department of Anatomy and Anthropology

Faculty of Medical & Health Sciences

Tel Aviv University

Ramat Aviv 6997801

Israel

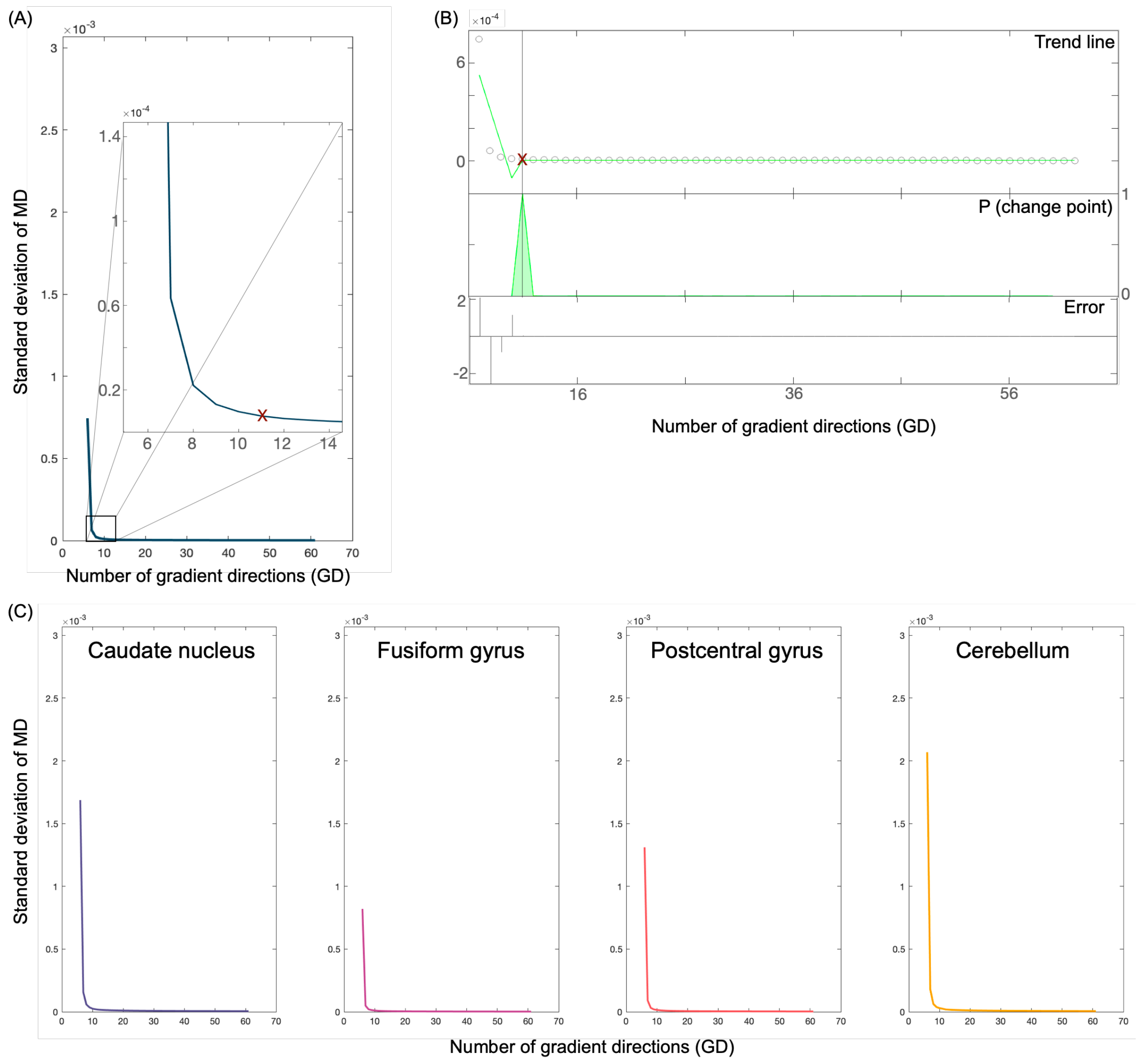


**Supplementary figure S1**. **MD stability for different window sizes based on excluded participants**. (A) SD of the mean MD in the gray matter calculated from an increasing number of gradient directions, starting with 6 and up to 61, based on 1000 permutations. The value for eleven gradient directions is marked by a red X. (B) Based on the BEAST algorithm (Zhao et al., 2019), we examined the changepoint of the curve after which the SD is close to zero, resulting in 11 gradient directions. (C) SD of the mean MD in four gray matter ROIs calculated from an increasing number of gradient directions, starting with 6 and up to 61, based on 1000 permutations. Based on four participants who were excluded from our study due to technical issues during the scan or excessive movement.

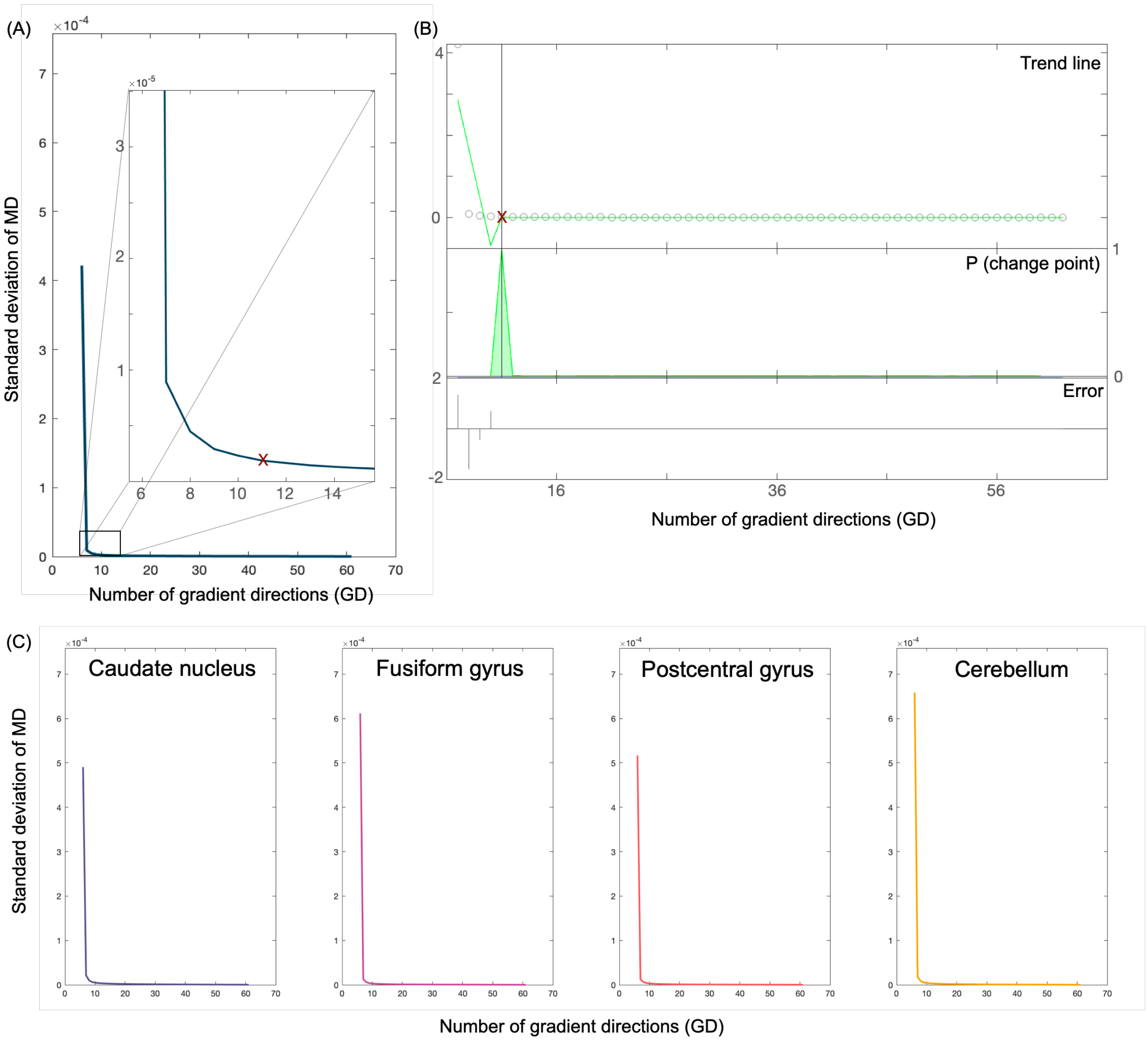


**Supplementary figure S2. MD stability for different window sizes based on external participants.** (A) SD of the mean MD in the gray matter calculated from an increasing number of gradient directions, starting with 6 and up to 61, based on 1000 permutations. The value for eleven gradient directions is marked by a red X.(B) Based on the BEAST algorithm (Zhao et al., 2019), we examined the changepoint of the curve after which the SD is close to zero, resulting in 11 gradient directions. (C) SD of the mean MD in four gray matter ROIs calculated from an increasing number of gradient directions, starting with 6 and up to 61, based on 1000 permutations. Based on ten participant who were scanned in the same scanner as our participants but using a different dMRI protocol.

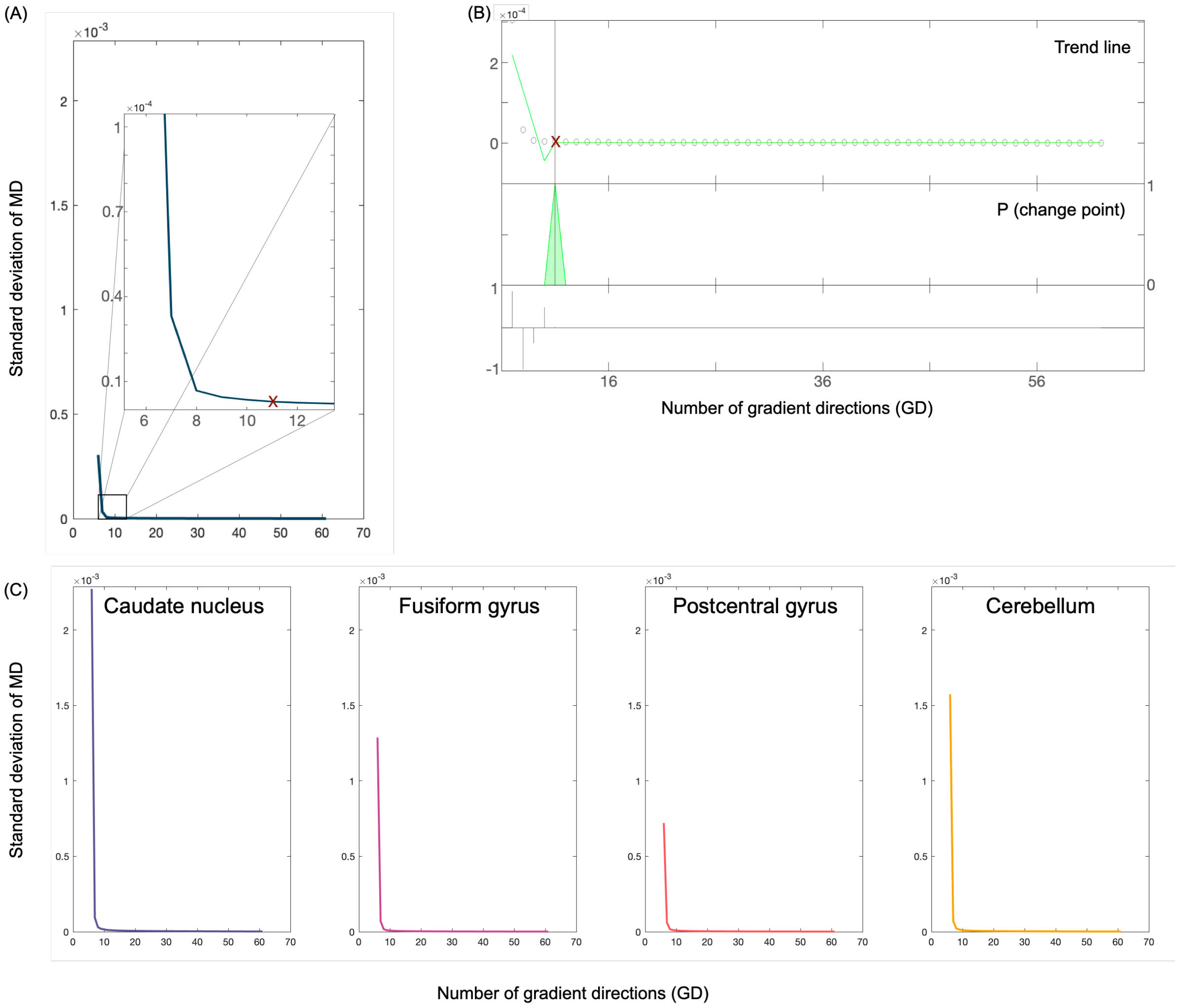


**Supplementary figure S3**. **MD stability for different window sizes based on HCP participants.** (A) SD of the mean MD in the gray matter calculated from an increasing number of gradient directions, starting with 6 and up to 61, based on 1000 permutations. The value for eleven gradient directions is marked by a red X. (B) Based on the BEAST algorithm (Zhao et al., 2019), we examined the changepoint of the curve after which the SD is close to zero, resulting in 11 gradient directions. (C) SD of the mean MD in four gray matter ROIs calculated from an increasing number of gradient directions, starting with 6 and up to 61, based on 1000 permutations. Based on five participants from the HCP database (Van Essen et al., 2012).

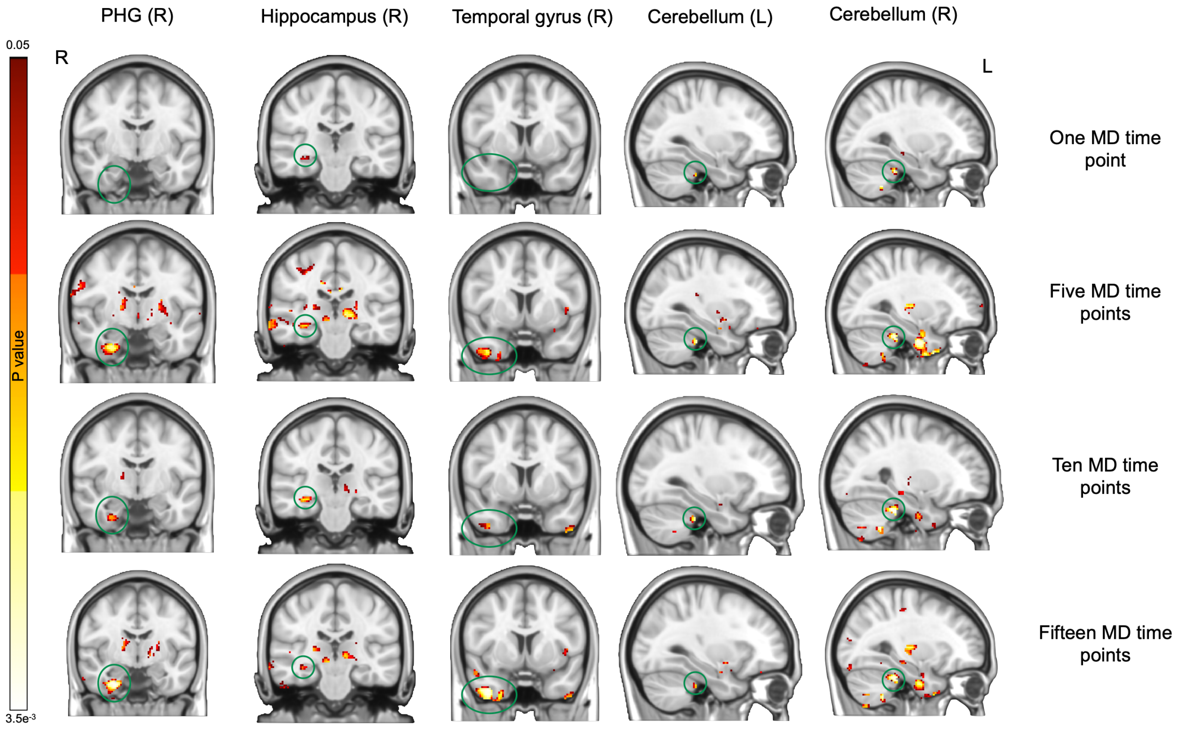


**Supplementary figure S4. Microstructural neuroplasticity at discrete timepoints throughout motor sequence learning based on different timepoints average.** The pre/post learning effect calculated from a 2 (learning vs. control groups) by 2 (pre/post) ANOVA. In each row, the pre- and post-learning MD is calculated as the average across different number of MD timepoints: the first and last timepoints (top row), the first and last five timepoints (second row), the first and last ten timepoints (third row) and the first and last fifteen timepoints (bottom row). Clusters used in the main text analyses are marked by a green circle. While not all clusters were detected when using only a single timepoint as the pre- or post-learning MD, they were found in all other options.

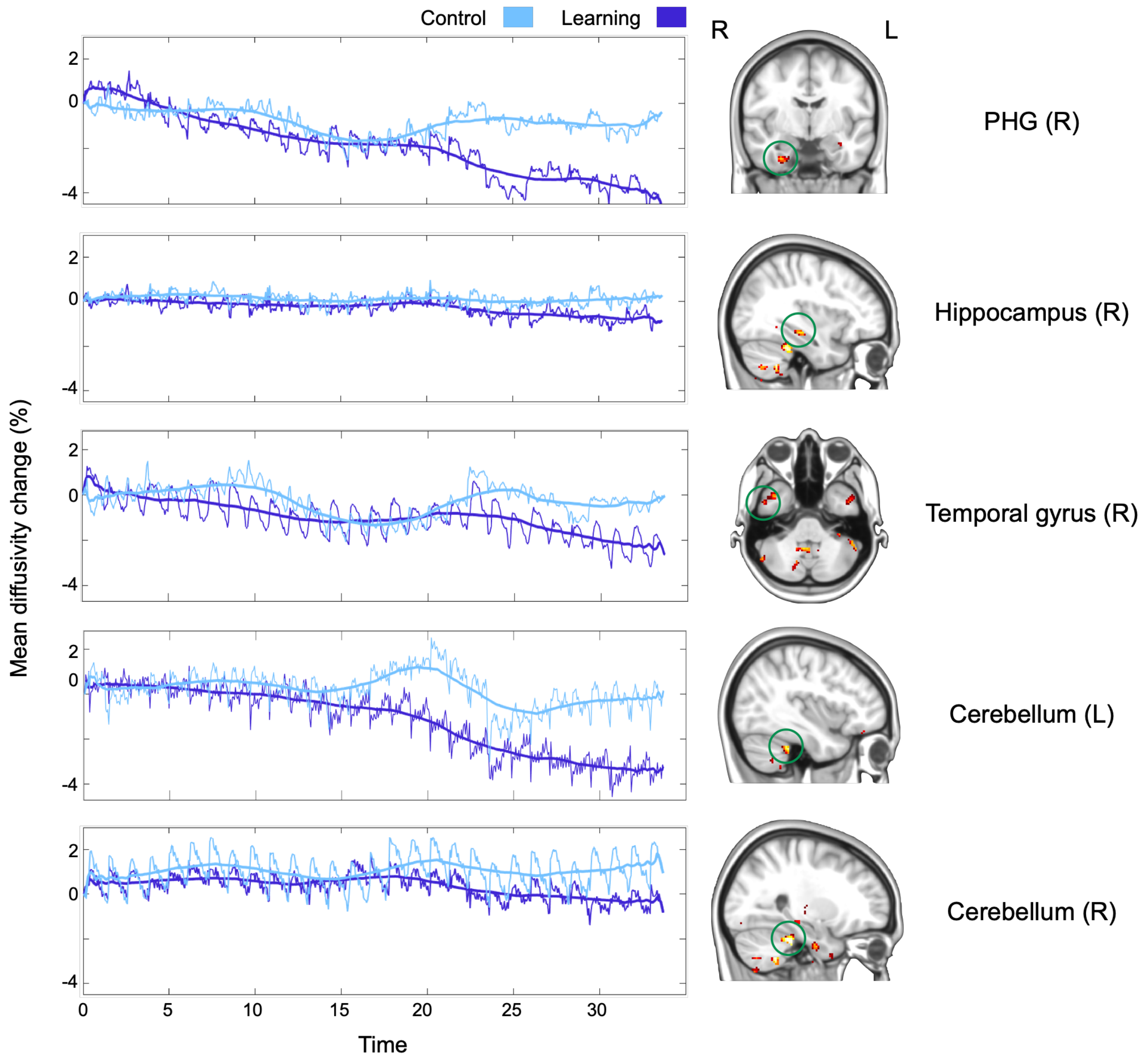


**Supplementary figure S5. Unfiltered continuous MD changes during learning.** Raw MD timeseries per cluster (before applying lowpass filter). Curves are calculated individually and averaged across participants per group. A 100-points smoothed curve is displayed to represent the trend of MD reduction during learning, averaged within group.

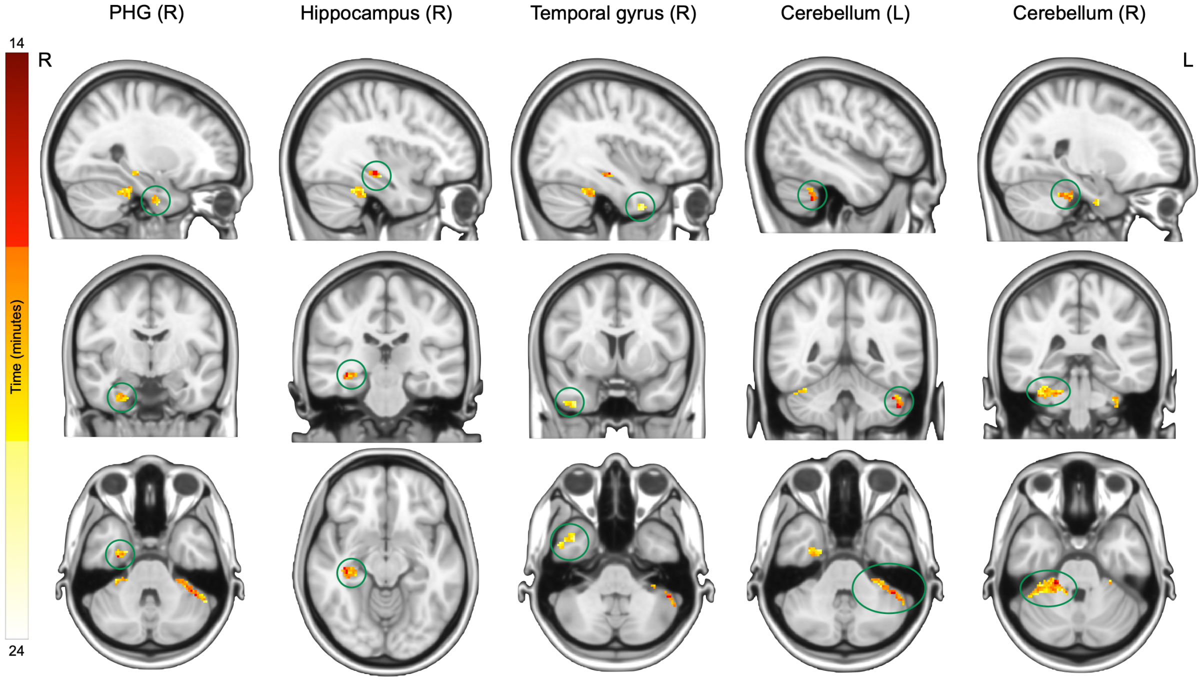


**Supplementary figure S6. Spatial changepoint detection analysis.** Voxelwise change-point detection within each of the five clusters for the task group. The change point was determined as the maximum change between the slope of the two parts of the divided curve.

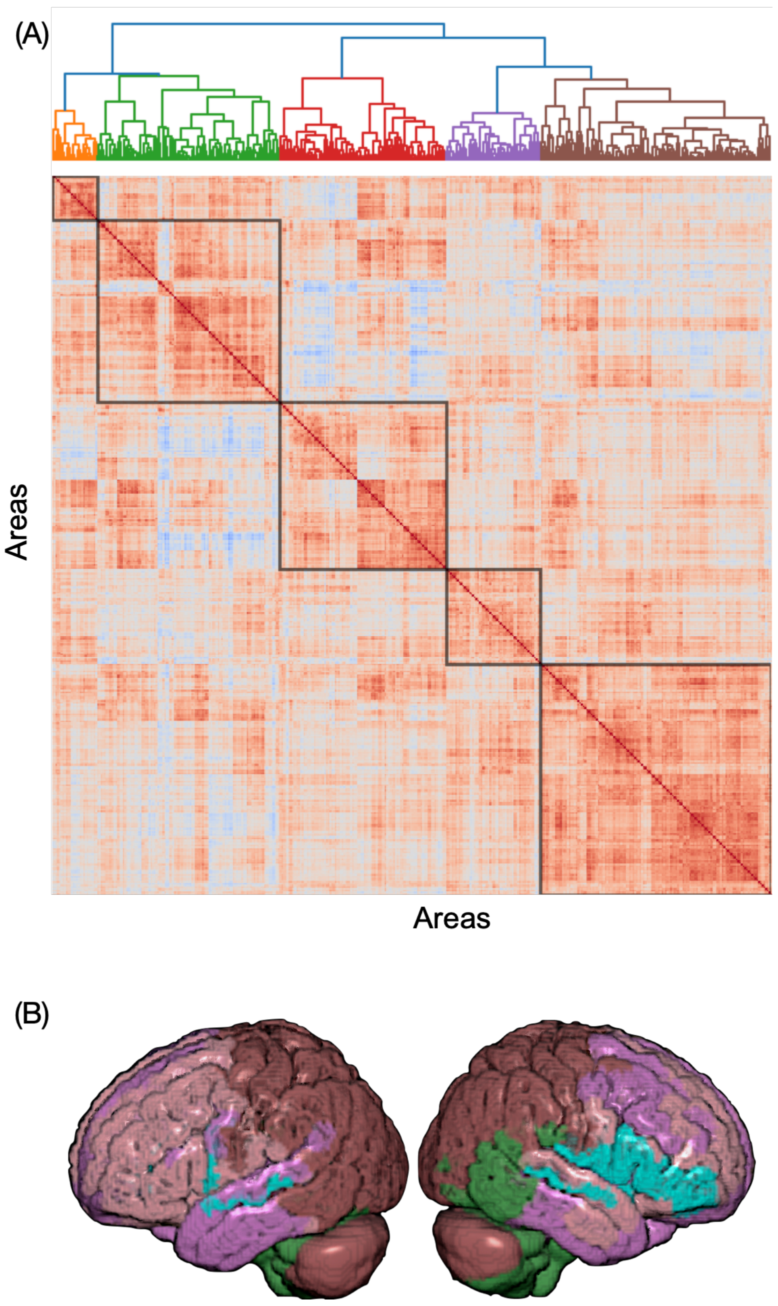


**Supplementary figure S7. Neuroplasticity networks originated from the Glasser atlas. (A)** A matrix depicting the pairwise Pearson's correlations between the patterns of MD decrease across 418 brain regions (parcellation of cortex and subcortex based on the Glasser atlas, (Glasser et al., 2016)), ordered by a hierarchical clustering algorithm and divided to neuroplasticity networks (black squares). **(B)** The five networks displayed on an inflated surface of the brain.

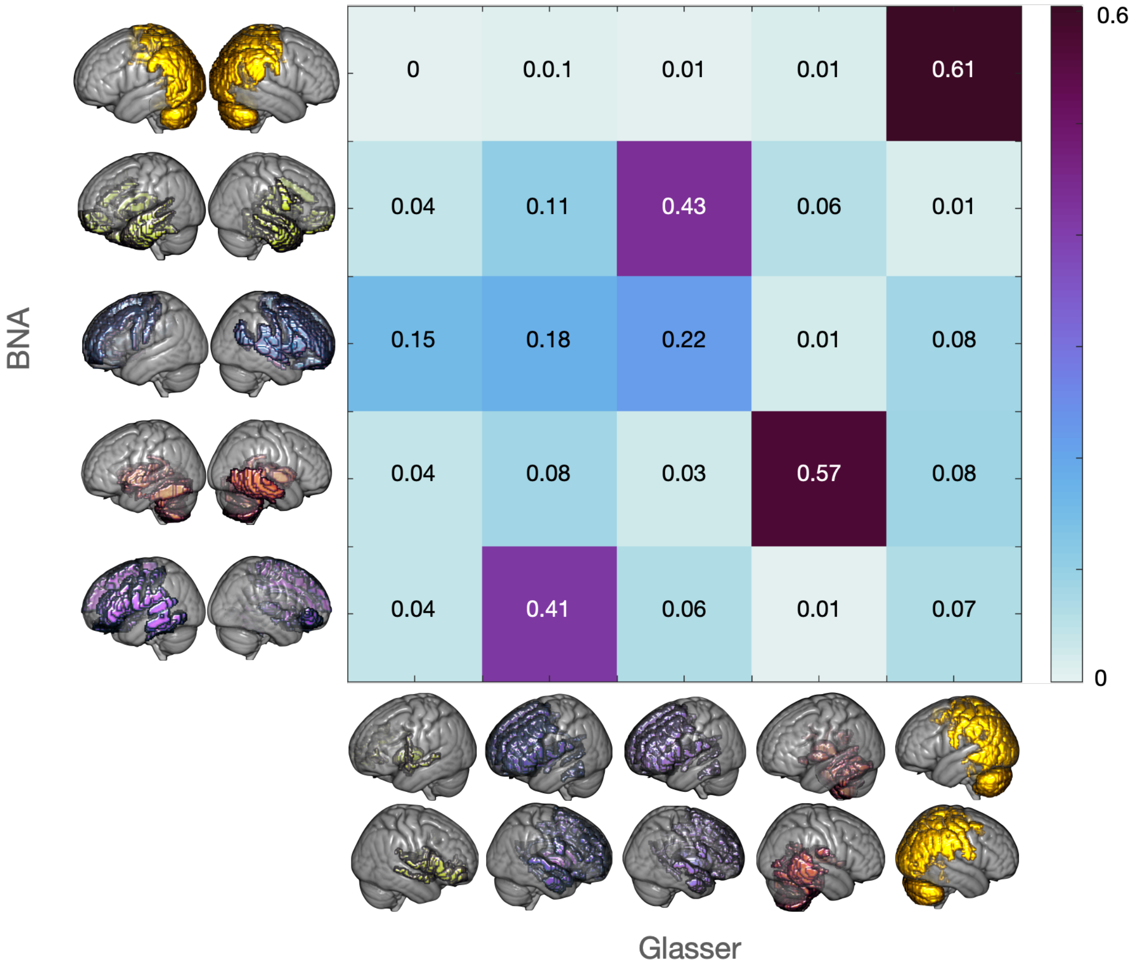


**Supplementary figure S8. Similarities between the Neuroplasticity networks originated from the Glasser atlas and the neuroplasticity networks originated from the BNA atlas.** Similarity matrix showing the Dice indices between each of the five neuroplasticity networks detected in the main analysis (based on the BNA parcellation, see Figure 6 in the main text) and the five networks based on the Glasser atlas (Supplementary Figure S7).

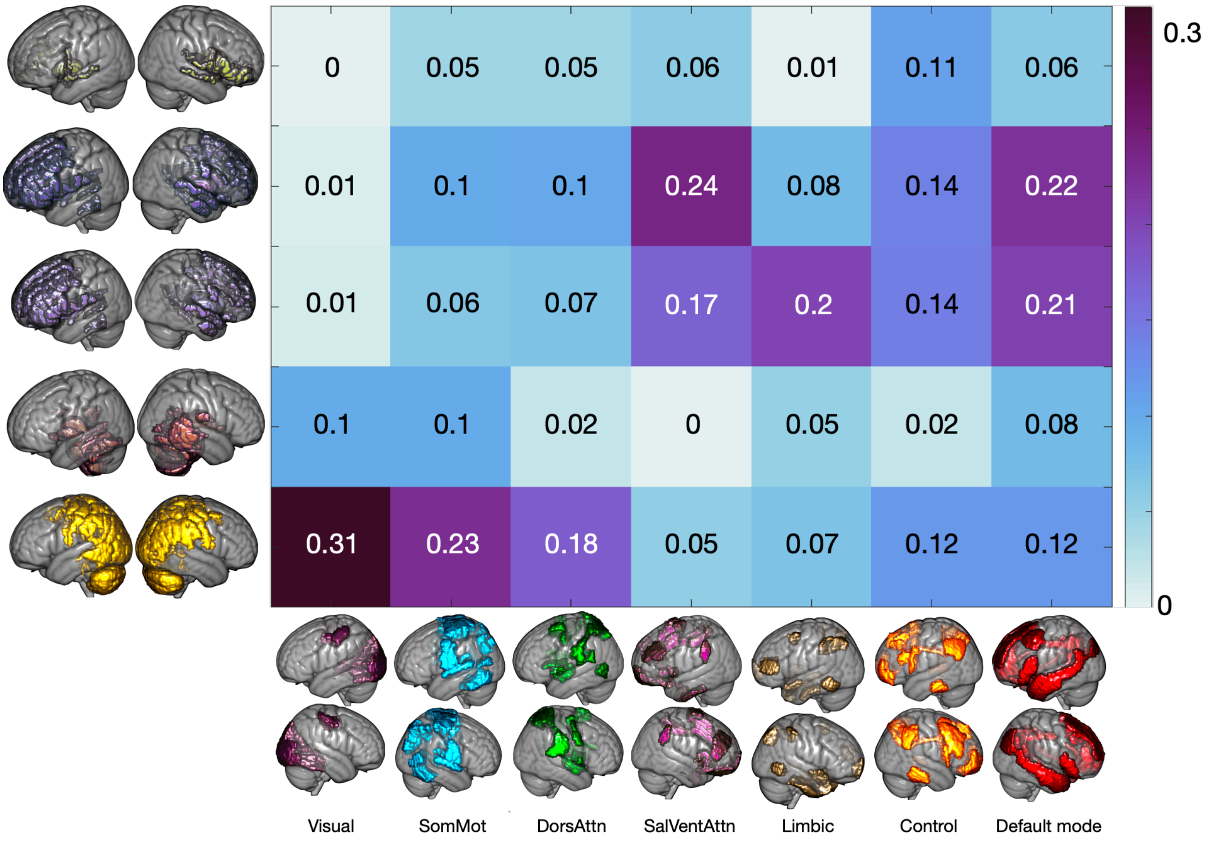


**Supplementary figure S9. Similarities between neuroplasticity networks originated from the Glasser atlas and functional connectivity networks.** Similarity matrix showing the Dice indices between each of the five neuroplasticity networks reported Supplementary Figure S7 and Yeo's seven functional connectivity networks.

2. Van Essen, D.C., Ugurbil, K., Auerbach, E., Barch, D., Behrens, T.E.J., Bucholz, R., Chang, A., Chen, L., Corbetta, M., Curtiss, S.W., Della Penna, S., Feinberg, D., Glasser, M.F., Harel, N., Heath, A.C., Larson-Prior, L., Marcus, D., Michalareas, G., Moeller, S., Oostenveld, R., Petersen, S.E., Prior, F., Schlaggar, B.L., Smith, S.M., Snyder, A.Z., Xu, J., Yacoub, E., 2012. The Human Connectome Project: A data acquisition perspective. Neuroimage.
